# Distinct Implicit Contributions to Action Selection and Action Execution in Sensorimotor Adaptation

**DOI:** 10.1101/2025.09.12.675892

**Authors:** Tianhe Wang, Tony Lam, Jordan A. Taylor, Richard B. Ivry

## Abstract

The sensorimotor system is continuously adjusted to minimize error. Current theories assume that this adaptation process entails the operation of multiple learning systems, with a key division between implicit and explicit components. Recent studies have revealed several inconsistencies regarding the characteristics and constraints of the implicit system, suggesting that the current framework is incomplete. Here, we propose that these conflicting findings can be understood by recognizing that there are multiple implicit subcomponents, distinguished by their distinct computational goals. One well-studied component is implicit recalibration, a process critical for action execution which uses sensory-prediction errors to automatically refine the sensorimotor map. Here we describe a second, novel component, implicit aiming, a process which contributes to action selection to achieve the specific goals. Through a series of studies, we find compelling evidence that those two implicit processes show a clear separation in their temporal stabilities and contextual modulations. These distinct properties correspond to different computational frameworks attributing learning dynamics to either contextual inference or cancellation of competing neural populations, respectively. Together, these findings suggest an alternative framework for sensorimotor adaptation based on the computational goals of the system rather than phenomenology.

## Introduction

Sensorimotor adaptation continuously recalibrates the mapping between motor commands and movement outcomes to compensate for unexpected errors induced by external (e.g., wind direction) or internal factors (e.g., fatigue)^1^. Research over the past two decades has made clear that this process can reflect the operation of multiple processes^2–6^. A primary distinction has been made between an explicit component that involves the use of conscious aiming strategies and an implicit component that involves a recalibration process which operates outside conscious awareness^2,7^. The distinction between explicit and implicit components is motivated by studies showing that these processes are sensitive to different error signals, operate on distinct timescales, and, to some degree, depend on the integrity of different neural regions^2,8–10^.

This first-level distinction does not imply that there is a unitary learning process within each branch of the hierarchy. Indeed, various lines of evidence suggest that there may be multiple forms of explicit^11,12^ and implicit learning^13^. For example, recent work has revealed that an explicit strategy can be implemented by an algorithmic process or retrieved from memory, each yielding different capacities and constraints^11^. Likewise, implicit learning has also been shown to entail several distinct processes. First, implicit learning seems to operate on (at least) two different time scales, one that is fast and volatile, and another that is relatively slower and stable^13,14^. Second, while sensory prediction errors were originally considered to be the sole driver of implicit learning^4,6,8^, emerging evidence suggests that task-level performance, and potentially reward, may also modulate implicit learning^15–17^. Third, future actions are implicitly biased toward recently performed actions, regardless of error-related feedback ^18–20^, a form of learning that has been attributed to an implicit Hebbian-like, use-dependent process.

While these features suggest potential subdivisions within the domain of implicit learning, several inconsistent findings have complicated attempts to characterize these processes and neatly accommodate them within the explicit-implicit dichotomy. For example, in visuomotor rotation tasks in which participants are unaware of the perturbation, implicit learning decays by 50% following just a one-minute break ^13,14^. In contrast, implicit learning, when elicited in response to task irrelevant feedback appears to remain stable over time, with minimal decay observed even over many days^21,22^. Similar inconsistencies are observed concerning the influence of contextual factors on implicit learning. When explicit contributions to learning are eliminated, contextual effects such as savings or spontaneous recovery are absent ^23,24^; in contrast, savings is observed when explicit processes are suppressed through a forced-response-time paradigm^13,25^.

While these observations point to the need for a more complex hierarchy of the processes contributing to sensorimotor adaptation, it is useful to consider an alternative taxonomic organization. Rather than start from phenomenology, we propose that the primary division should be based on the computational goals of the system. In particular, we propose an initial division between processes involved in action selection and those involved in action execution (Fig. 1). The former would include processes involved in identifying the optimal action to achieve a goal; the latter would be processes involved in ensuring the precise implementation of the chosen action.

**Fig. 1.**
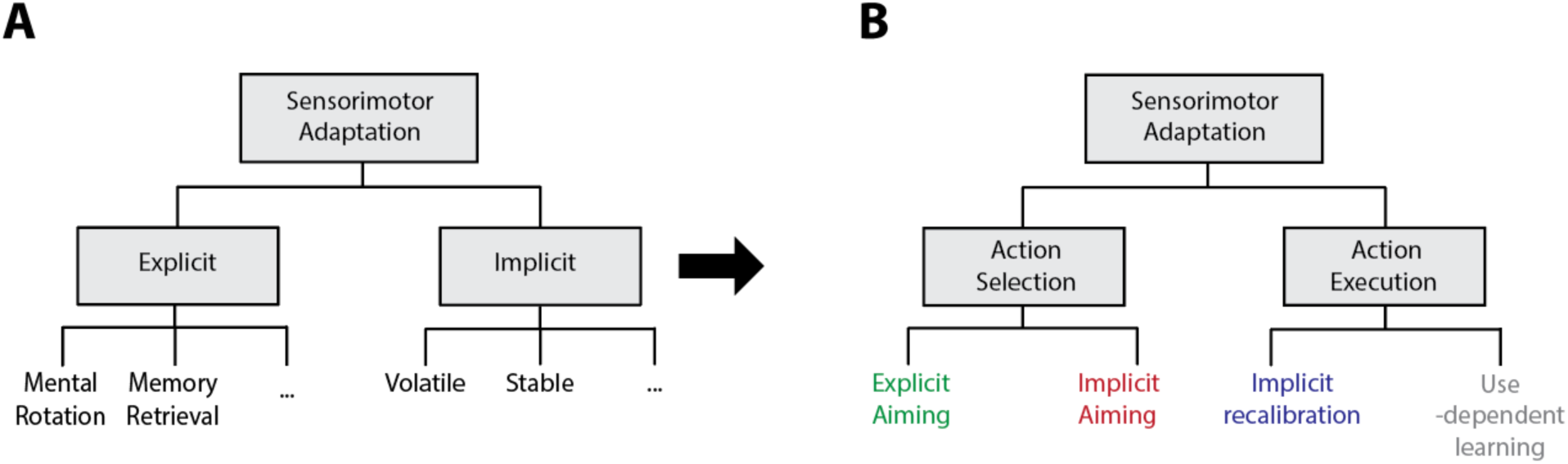
An alternative taxonomy of sensorimotor adaptation. (A) The classic taxonomy makes a primary distinction between implicit and explicit components, each of which may include multiple processes (not shown). (B) In an alternative taxonomy, the top branch is defined by the computational goals. Action selection includes processes that compute the action required to achieve the task goal. Action execution includes processes that ensure that the selected movement is accurately implemented. The execution system is hypothesized to include only implicit processes, whereas the selection system includes implicit and explicit processes.

By this taxonomy, action execution might be intrinsically implicit, encapsulated from the output of explicit processes (Fig. 1). For example, implicit recalibration responds in an obligatory manner to adjust the sensorimotor mapping based on sensory prediction errors even when its operation is sub-optimal in terms of achieving task success^26,27^. Similarly, the bias to repeat a motor command that underlies implicit use-dependent learning imposes biases towards recently executed movements even when the target location has changed ^18,28^.

In contrast, action selection may reflect the operation of both explicit and implicit processes. Clearly the utilization of a strategy can be intentional and explicit; for example, the participant might make a conscious decision to modify an action plan to counteract a perturbation. But there is also evidence to suggest that action selection can be influenced by unintentional and implicit processes. For example, there may be unconscious contributions to aiming dependent on recent history^20^. Indeed, a large body of work has shown how habits influence choice behavior outside conscious control^29–31^, a manifestation of use-dependent learning at the selection level. Moreover, previous studies demonstrated that when participants adjusted their aiming direction without awareness in response to a shifting reward landscape^32–34^.

In the present study, we conducted a series of experiments to explore implicit contributions to action selection, and in particular, a process we will refer to as implicit aiming. We employed a variety of methods to identify and characterize this process, highlighting how it is distinct from implicit recalibration. The results show that these two forms of implicit learning differ in their temporal dynamics and sensitivity to contextual information. When considering the nature of implicit and explicit processes to action selection, we hypothesize that a core difference, at least in sensorimotor adaptation tasks, may arise from a credit assignment process. Explicit re-aiming is invoked when errors in performance are attributed to an external perturbation; in contrast, implicit aiming is invoked when errors in performance are attributed to internal motor biases or proprioceptive drift^35,36^.

## Results

### Experiment 1: Implicit aiming in visuomotor adaptation

Exp. 1 was designed to identify an implicit process that contributes to action selection. In the main phase of the experiment, participants performed a visuomotor rotation task. Feedback was limited to a cursor that appeared after the movement, with the position of the cursor contingent on the position of the hand and modified by the current size of the perturbation. During the rotation phase, participants were instructed that their task was to hit the target with the cursor. The task was designed to measure performance when the contributions of implicit recalibration and explicit aiming were restricted. For the former, we delayed the endpoint feedback by 1.5 s, a manipulation that has been shown to essentially abolish implicit recalibration^37,38^. For the latter, we increased the perturbation in a gradual manner, increasing the angular perturbation by 0.5°/trial up to a maximum value of 65°. While it is likely that at some point, participants become aware that the visual feedback is perturbed, the uncertainty introduced by the gradual change limits the utility of a specific aiming strategy^32,39,40^. To further increase the uncertainty of the perturbation size we gradually decreased the rotation from 65° to 25° over the second half of training.

To assess the contribution of an implicit component to the observed changes in behavior, we inserted probe blocks (P1-P3) in which no visual feedback was provided, and participants were instructed to move directly to the target^7,9^ (Fig. 2A). We attribute systematic deviations in reaching direction on these trials to implicit aiming. We note that, at this stage, implicit aiming could theoretically reflect use-dependent learning, reward-based learning, or some other learning processes. We will address this issue in subsequent experiments.

**Fig. 2.**
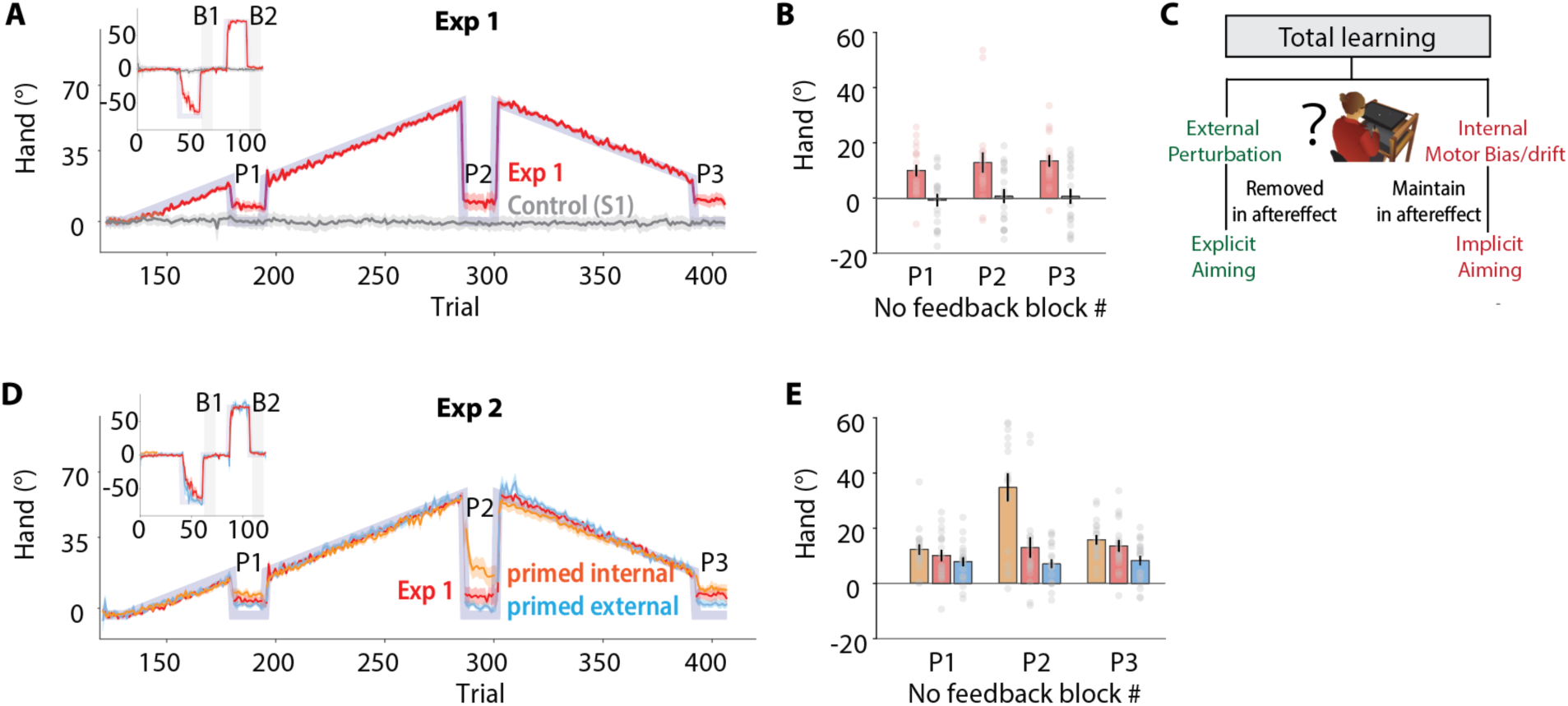
Implicit aiming during sensorimotor adaptation with delayed feedback. (A) Exp. 1: The perturbation schedule (transparent purple) and mean reach angle for the groups receiving contingent feedback in the main experiment (red) or non-contingent, clamped feedback in the control experiment S1 (grey). Feedback was delayed for 1.5 s in both conditions to attenuate implicit recalibration. P1 - P3 refer to the three probe blocks in which reach direction was measured in the absence of visual feedback and participants were instructed to reach directly to the target. Inset shows perturbation and reach angle during a baseline phase that consisted of blocks of veridical feedback, rotated feedback that was introduced in an abrupt manner (70°) and two no-feedback blocks (B1 and B2). (B) Mean reach angles in the three no-feedback blocks of Exp. 1. (C) Illustration depicting the credit assignment problem that arises at the start of each probe block. Participants will adjust their reach angle based on their estimate of how much of the change in performance was due to their use of a strategy to offset the external perturbation. The aftereffect reflects implicit changes in performance that arise from internal motor biases and/or proprioceptive drift. (D) Exp. 2: Time course of hand angle in Exp. 2 for group primed for the external perturbation (blue) and group primed to anticipate motor bias (yellow). For comparison, the data from Exp. 1 are replotted (red). (E) Aftereffect in the three no-feedback blocks (E1-E3). Shaded areas in A, B, E and error bars in C, F indicate standard error.

To ensure that the participants understood the instructions on probe trials, namely, to aim directly towards the target, we included two manipulation checks before the main phase of the experiment. In each of these, we briefly introduced a 65° rotation for 20 trials. Participants were informed about the nature of the perturbation and instructed to use an aiming strategy to compensate for it. These short blocks were immediately followed by probe trials. Participants successfully adapted to the 65° abrupt rotation and showed minimal aftereffects (t(18)s<1.7; *p*s>0.1; Fig. 2A, intercept), demonstrating that they were able to adopt and then suppress an aiming strategy. Given these results, we assume that the participants understood the instructions to stop using explicit aiming strategies on probe trials.

During the main phase of the experiment, participants effectively adjusted their reach direction to offset the gradual rotation (Fig. 2A). The change in hand angle closely mirrored the perturbation function, indicating that the endpoint cursor landed near the target throughout the adaptation phase. Note that this learning could include both implicit and explicit processes.

Turning to the results of primary interest, we observed a pronounced aftereffect in all three probe blocks (Fig. 2B). While there was a large drop in heading angle with the onset of each of these blocks, the heading angle remained biased in the direction opposite of the target (P1: 10.0 ± 2.1°, P2: 13.0 ± 3.7°; P3: 13.6 ± 2.1°, t(18)s>3.5, *p*s<.002). Interestingly, the magnitude of this effect was considerably larger than the aftereffects observed on the manipulation check blocks in which the probe trials were presented after an abrupt perturbation (t(18)s>2.7, *p*s<.01). We also conducted two control experiments, confirming that these aftereffects were not due to implicit recalibration or use-dependent learning (See Supplementary Results 1-2).

### Experiment 2: Implicit aiming arises from a credit assignment problem

Having identified a form of implicit learning distinct from implicit recalibration and use-dependent learning, we next sought to characterize the source of this process. We considered three hypotheses. First, implicit aiming could reflect the outcome of a process in which an explicit aiming strategy becomes automatized, a type of habit formation. Second, implicit aiming may reflect a reward-based learning process. Implicit shifts in reach direction are observed in response to a gradual perturbation when binary feedback is provided to indicate success or failure^32–34^. The effect of the cursor hitting or missing the target during rotation training in Exp. 1 could have served as a reward signal driving an implicit reward-based learning process.

Third, the aftereffects in Exp. 1 may arise from confusion created in attributing the observed error to a change in the external environment or internal motor biases^41^. The former would correspond to the participant’s estimate of the experimenter-imposed perturbation; the latter would correspond to the participant’s sense that some of the error is due to idiosyncratic motor biases^42,43^ and proprioceptive drift that occurs in the absence of vertical feedback^35,44^. In the probe trials, we expect they would stop compensating for the former component, reducing their aim by the estimated external perturbation. However, the latter should continue to be relevant with the size of the aftereffect on the probe trials providing an estimate of this component. The absence of an aftereffect in the two baseline probe blocks of Exp. 1 indicate that solving this credit assignment problem might be simple for a large, invariant perturbation that is introduced abruptly. But it could be challenging in conditions where the perturbation changes continuously. Here, participants may attribute part of the external perturbation to internal motor biases, an attribution error that would be manifest as a persistent aftereffect.

To empirically evaluate these three hypotheses in Exp. 2, we manipulated the awareness of the perturbation by providing different “priming” instructions to two groups prior to the onset of the gradual rotation. For one group, we described the forthcoming rotation schedule in detail, including the way in which the size of the perturbation was incremented in small steps until reaching a maximum value of 65°. The other group was simply told that people tend to become biased in the direction of a repeated movement over time, especially when they cannot see their hand. In this way, we primed the first group to attribute the error to the external perturbation (external-bias group) and primed the second group to anticipate a motor bias (internal-bias group). We also removed the abrupt rotation manipulation check for the internal-bias group to avoid priming those participants to attribute subsequent errors to an external perturbation.

The hypotheses that explicit aiming transitions into an implicit process or that implicit aiming reflects a reward-based learning process predict that the priming instructions should have no effect on the aftereffect: The reward signal and the aiming direction during training are similar across the two conditions. However, based on the credit assignment hypothesis, we predicted that priming would have opposite effects on the two groups. The external prime should reduce the aftereffect since these participants now anticipate the rotation and, thus, are more likely to be sensitive to their aiming strategy. In contrast, the internal prime should result in an increase in the size of the aftereffect given that these participants will assume some part of the change in heading direction is due to an internal motor bias.

Both groups of participants in Exp. 2 performed well in compensating for the gradual perturbation and the learning curves closely mirrored that in Exp. 1. Moreover, both groups showed significant aftereffects in the no-feedback blocks (Fig. 2E, ts>4.8, *p*s<.001). Consistent with the prediction of the credit assignment hypothesis, the size of the aftereffect was much larger in the internal-bias group compared to the external-bias group, even though both groups experienced the same gradual perturbation (*F*(166)=5.2, *p*<.001, Fig. 2F). Moreover, when compared to the results of Exp. 1, priming an internal bias increased the aftereffect (change from E1: 6.8 ± 2.2°, *t*=3.1, *p*=.002), whereas priming an external bias decreased the aftereffect (-4.4 ± 2.1°, *t*=-2.0, *p*=.041). The difference between conditions was most prominent in the second no-feedback block when the perturbation was at a maximum. These results are at odds with the reward and habitualization hypotheses as both accounts predicted no effect of the priming instructions.

### Experiment 3: Implicit aiming shows saving whereas the implicit recalibration shows anti-saving

Having established an implicit process that impacts action selection, we next sought to characterize its dynamics in comparison to implicit recalibration, the implicit process that impacts action execution. In Exp 3 we ask if implicit aiming exhibits savings, the phenomenon in which learning is facilitated upon re-exposure to a previously experienced perturbation. Prior work has convincingly shown that, in standard visuomotor adaptation tasks, savings is associated with the explicit component of action selection: Having discovered an appropriate aiming strategy when first encountering a perturbation, participants can readily recall that strategy when the same perturbation is reintroduced after an extended washout period^45^. In contrast, implicit recalibration does not exhibit savings. Indeed, when learning is restricted to implicit recalibration with clamped feedback, savings is attenuated^24,46^.

However, as noted in the Introduction, the picture is muddied when considering the results from other experimental manipulations designed to isolate implicit learning. For example, savings is observed when planning time – and thus explicit contributions to aiming – is minimized via a forced reaction time task^13,25,47^. We hypothesize that this discrepancy arises because the paradigms tap into different forms of implicit learning. The clamped feedback procedure isolates implicit recalibration whereas the forced response time task also allows for the operation of implicit aiming. By this view, we hypothesize that implicit aiming will exhibit savings.

To test this hypothesis, we used the forced response time task and compared the degree of observed savings following delayed contingent feedback versus clamped non-contingent feedback. The experiment consisted of two rotation blocks, each with a 30° abrupt rotation, separated by a no-feedback block to assay aftereffects and a feedback block to fully wash out residual learning from the first rotation block. On each trial, participants heard a countdown of four tones (separated by 500 ms) and were required to initiate their movement with the final tone^48^. Critically, the target appeared at one of three locations 250 ms prior to the last tone, leaving insufficient time to implement an explicit aiming strategy^11,25,48^. Given that learning under such conditions is primarily dependent on implicit processes, we assumed that the delayed contingent feedback would isolate implicit aiming and that the clamped non-contingent feedback would isolate implicit recalibration. Note, because participants are instructed to aim directly to the target and ignore the cursor feedback in the clamp condition, we assume that there should not be any aiming, explicit or implicit.

In both conditions, reaction times averaged around 220 ms throughout the whole experiment (Fig. 3B). This RT is similar to the values observed in forced RT reaching tasks with multiple targets^11,25,48^, consistent with the assumption that the contribution of explicit aiming was negligible. Importantly, we observed a gradual change in reach angle in response to the delayed feedback (Fig. 3D), suggesting that implicit aiming operates under limited RT conditions.

**Fig. 3.**
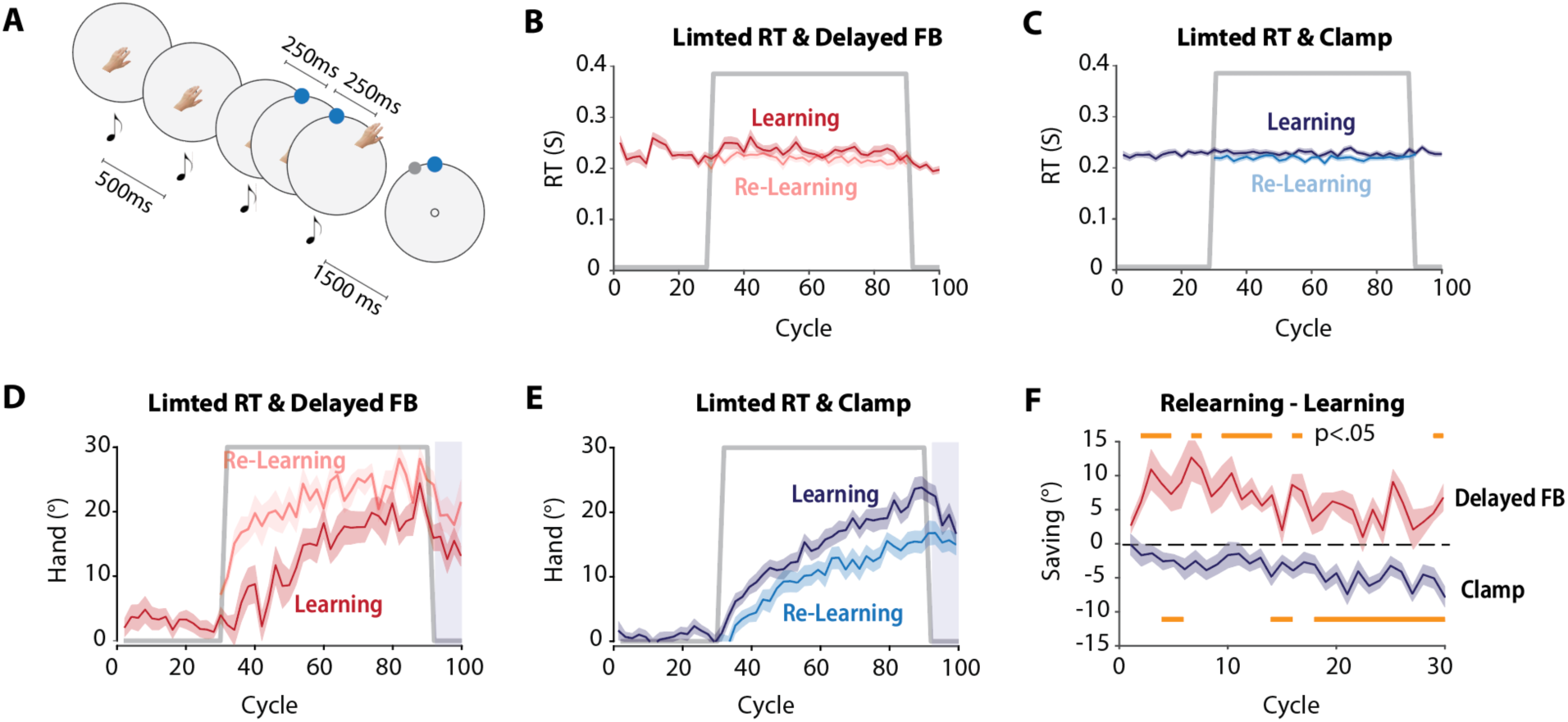
Implicit aiming shows savings, while recalibration shows anti-savings. (A) Forced RT task to minimize explicit aiming. The target appears 250 ms after the third tone, and participants are instructed to initiate their movement with the fourth tone. In separate groups, we used delayed contingent feedback or non-delayed clamped feedback. For the former, the rotated feedback cursor appeared 1.5 s after the hand reached the target distance; for the latter, the clamped (noncontingent) feedback cursor appeared as soon as the hand reached the target distance. (B–C) Average reaction times were around 220 ms and were similar in the two perturbation blocks. (D) In response to the contingent rotation, participants exhibited savings during relearning. The aftereffect data indicate that learning is mostly implicit aiming (>10d). The gray line represents the perturbation schedule. (E) In response to clamped feedback, participants exhibited attenuation during relearning. (F) Difference between the first (learning) and second (re-learning) perturbation blocks. The yellow bars above and below indicate epochs in which the values for their associated function are significantly different from zero.

While the learning functions in the two conditions were quite similar in response to the initial perturbation, there was a marked divergence during the relearning phases. Consistent with prior work^23,24^, learning was attenuated in the clamped feedback group. In contrast, a strong savings effect was seen in the contingent feedback group. A mixed ANOVA showed a significant interaction effect between feedback type and block number (p<.0001). As a *post-hoc* test, we employed a cluster analysis to compare learning and relearning for each feedback condition. We observed significant attenuation during late training for the clamped feedback group and significant savings during early training for the delayed feedback group (See Fig. 5F).

### Experiment 4: Different temporal stability of implicit aiming and implicit recalibration

The temporal dynamics of implicit adaptation from a single trial has also been the focus of considerable interest. One prominent hypothesis is that implicit adaptation entails two components, one that is temporally volatile and the other that is temporally stable. This hypothesis builds on the observation that the adaptive change in heading angle to a visuomotor rotation can be reduced by up to 50% following a 1-minute break^13,14^. Moreover, the size of trial-by-trial change of hand angle in response to random perturbation is inversely related to the inter-trial-interval^23^. Here we asked if these two components are dissociable in terms of implicit aiming and implicit recalibration.

To examine the temporal dynamics of implicit aiming, we applied a delayed feedback task with three key manipulations (Fig. 4A). First, we applied an abrupt 30° rotation which was then gradually decreased to 16° over 100 trials. This gradual reduction was included to mimic how the error signal diminishes in classic visuomotor rotation tasks due to implicit recalibration. Second, we did not provide any feedback during the baseline phase, introducing the 30° rotation on the first trial with cursor feedback. We reasoned that, in the absence of veridical feedback, participants would be more likely to attribute the resulting error to their own motor bias rather than to an external perturbation. Third, and most importantly, we inserted three 1-minute breaks during the gradual rotation phase. The reduction in heading angle following these breaks provides a measure of temporal volatility (Fig. 4B). To compare implicit aiming and implicit recalibration, we tested a second group of participants with a similar design but used non-delayed clamp feedback (Fig. 4C-D).

**Fig. 4.**
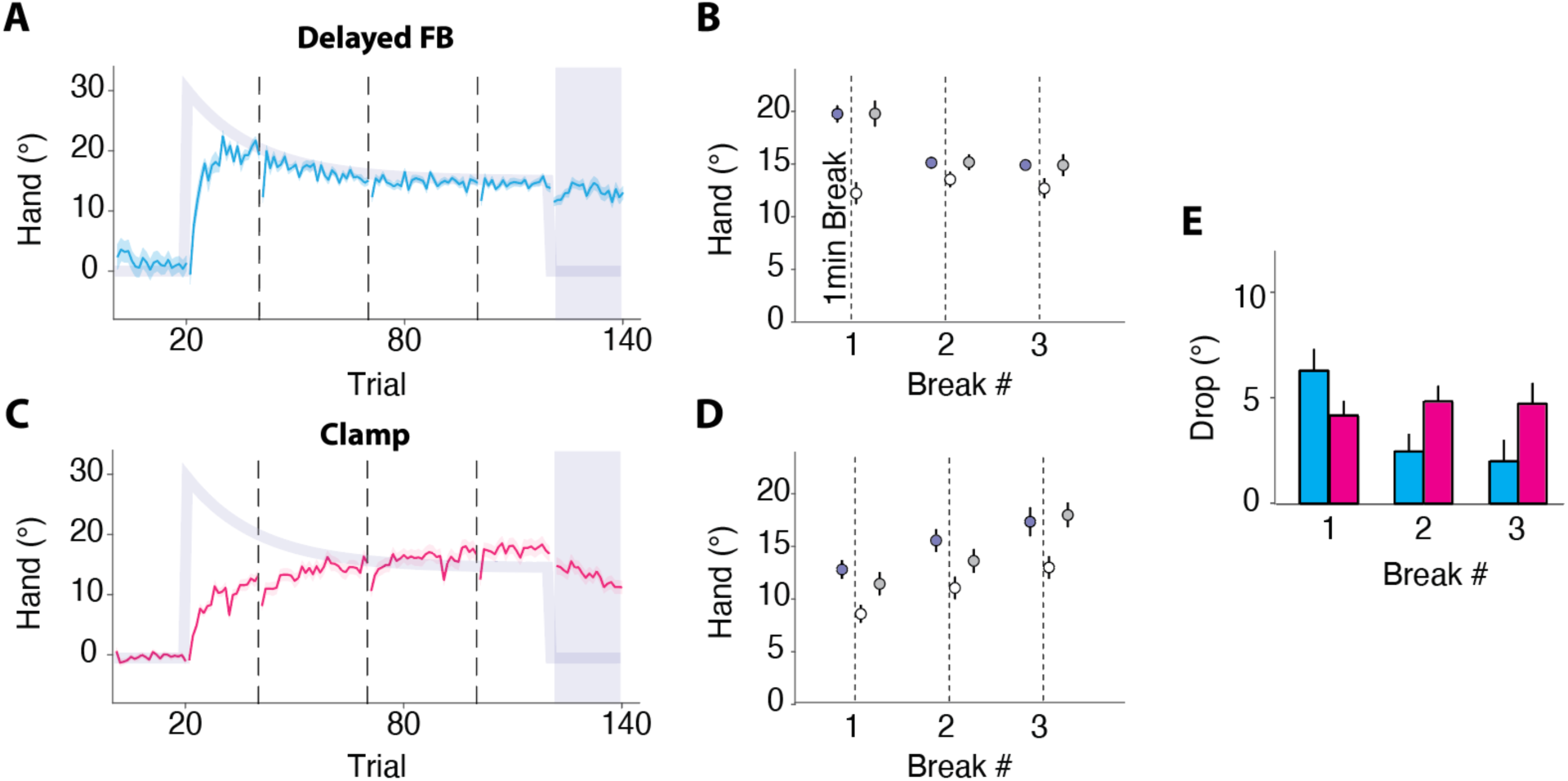
Temporal stability of implicit aiming. (A) Time course of hand angle in Exp 4a (delayed task-contingent feedback). Light purple line indicates perturbation schedule. Vertical dashed lines denote the three 1-min breaks. (B) Comparison of reach angle, before and after the breaks. Purple dots show averages of three pre-break trials. White and gray dots represent first and second post-break trials, respectively. Significant drops occurred after each break, with rapid recovery (presumably due to error feedback experienced on the first post-break trial). (C-D) Equivalent analyses to A-B for non-delayed clamped feedback group. (E) Reduction in hand angle for the delayed, contingent feedback group (blue) and non-delayed clamped feedback group (red).

Participants showed prominent learning in both conditions. In the delayed feedback condition, participants showed a rapid change in hand reach angle following the abrupt perturbation (Fig. 4A). On average, heading angle adjustments matched the perturbation size after 20 trials and effectively countered the declining rotation thereafter. The average heading angle in the aftereffect block was very similar to late perturbation performance (Fig. 4A, t(26) = 0.8, p=0.45), suggesting that learning here was almost all due to implicit aiming. We assume that absence of an initial veridical feedback block coupled with the relatively small size of the perturbation eliminated an explicit aiming component. Learning was slower for the clamped feedback group and reached a level that slightly surpassed the perturbation in late learning.

Our primary interest was on the behavioral change following the 1-minute breaks. In both conditions, performance drops were observed after each break (Fig. 4E, all p’s<0.05), the signature of a volatile contribution to learning. We note that if we sum the size of drop measured in both conditions, the combined drop is similar in magnitude to that observed in visuomotor rotation studies using this probe technique^13,14^. Thus, the current results suggest that there is a volatile component in both implicit aiming and implicit recalibration.

Notably, however, a comparison of the two conditions revealed that the pattern of the drop across the three probes differed for the two conditions (Mixed linear model, p(group x time) = 0.009). For the delayed contingent feedback condition, the reduction was largest after the first break (6.3°, 36%) and decreased after the other two breaks (∼3° or 20%, p = 0.001). For the non-delayed clamped feedback recalibration, the reduction remained relatively constant across the three probes (∼5°, p=0.6). This interaction suggests that the mechanisms for the post-break performance drops may differ for implicit aiming and implicit recalibration.

To explore this hypothesis, we considered two computational models that have been proposed to explain the dynamics of action selection and implicit recalibration. The Contextual Inference (COIN) model^49^ focuses on the dynamics of action selection by postulating that the motor system establishes unique motor memories for separate contexts. For example, the baseline phase (veridical feedback) and perturbation phase (rotated feedback) would define two contexts (Fig. 5A). Participants infer the current context to determine which memory to express. The Cerebellar Population Coding (CPC) model^23^ draws on the physiology of the cerebellum to account for the dynamics of implicit recalibration. Of primary relevance, recalibration arises from the population activity of units tuned to different directions in the cerebellar cortex and deep cerebellar nuclei (DCN) (Fig. 5C)^50,51^. In contrast to COIN, this model does not represent context; rather, learning dynamics emerge from shifts in the balance of activity among populations tuned to opposing directions.

**Fig. 5.**
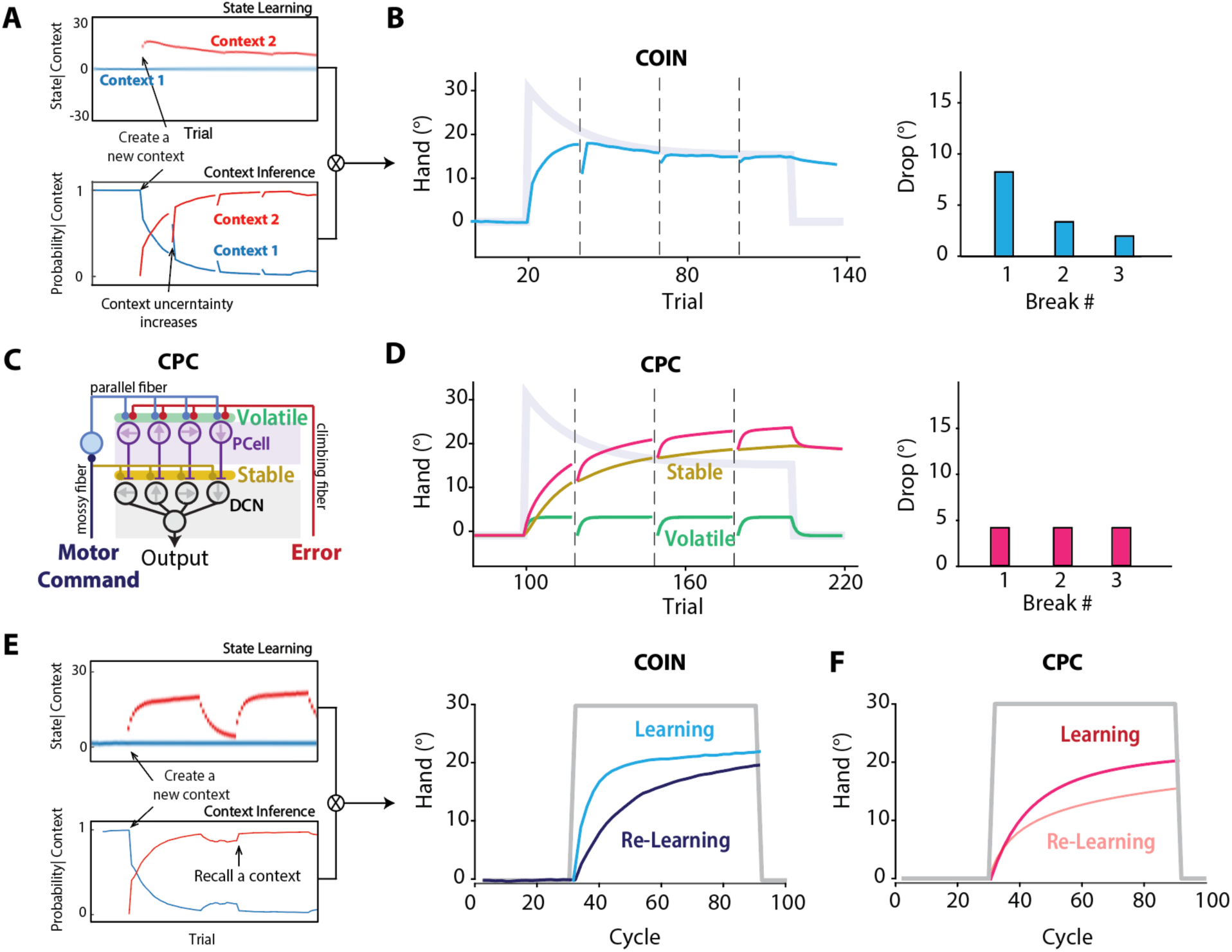
Distinct computational mechanisms explain implicit aiming and implicit recalibration properties through COIN and CPC models respectively. (A) In the COIN model, participants establish two contexts during training (baseline and perturbed contexts). Reach angle reflects a weighted combination of the state estimation for each context and their current relative probabilities. The COIN model predicts that context uncertainty increases during the 1-min breaks. As the number of training trials increases, context uncertainty decreases as the prior for the perturbed context increases. (B) As such, the drop after the 1-min break decreases across time. (C) The CPC model involves a two-layer learning system (cerebellar cortex and DCN) with different retention rates. (D) In the CPC model, total learning combines volatile (cortex) and stable (DCN) components. Drops after the 1-min breaks reflect forgetting in the volatile layer. The magnitude of this drop remains constant throughout training given that the learning and retention rates are fixed. (E) Savings during relearning is predicted by the COIN model. When the same perturbation is reintroduced, participants rapidly retrieve the learned context. (F) The CPC model produces attenuation during relearning (see Fig. S3 for mechanisms).

We simulated both models to evaluate their ability to explain the temporal dynamics in Exp 4. Both models predict declines in the reach angle following a 1-min break (Fig. 5B, D), although the mechanisms are different. In the COIN model, context uncertainty increases during the breaks given the lack of environmental input^52^. This leads to the expression of mixed memories post-break due to reduced confidence in the active context (Fig. 5A). As such, the COIN model attributes the drop in reach angle after the break to context uncertainty rather than memory decay. The CPC model attributes the drop to differential retention rates between its layers. Drawing on physiological recordings in eyeblink conditioning, we assume that plasticity in the cerebellar cortex is relatively volatile characterized by fast learning and forgetting.^53–55^ In contrast, plasticity units in the DCN is relatively stable, supporting stable memories (Fig. 5C).

Critically, the models differ in how the magnitude of the drop changes over the course of the experiment. The COIN model predicts a strong drop in early training when the expectation of the baseline context is high (Fig. 5B). As training continues, the expectation for the perturbation context increases and, thus, the drop becomes smaller. This matches the pattern observed in the delayed feedback condition in which we assume learning is due to implicit aiming (Fig. 4F). In contrast, the CPC model predicts a consistent drop size throughout training (Fig. 5D). This is because the population dynamics within the cerebellar cortex remain consistent across training due to the absence of any context representation (i.e., memory). This prediction aligns with the pattern observed in the clamped feedback condition in which we assume learning is due to implicit recalibration.

Notably, the two models also capture the opposing relearning effects for implicit aiming and implicit recalibration observed in Exp 3. The COIN model predicts savings (Fig 5E). When the perturbation is reintroduced, the system rapidly identifies the perturbation context and retrieves the associated response. In contrast, the CPC model predicts anti-savings (Fig 5F). Relearning is slower due to interference from residual activity in units tuned to the opposite error direction during washout (Fig. S3). This dissociation is consistent with the idea that COIN provides a reasonable model of implicit aiming while CPC provides a reasonable model of implicit recalibration. More generally, our simulation results highlight how implicit aiming and implicit recalibration operate under distinct computational principles.

A full comparison here would require an evaluation of COIN’s ability to predict learning in response to clamped feedback and the CPC’s ability to predict learning in response to delayed feedback. However, COIN cannot be meaningfully simulated with clamped feedback (see Methods). The model relies on inferring the context and perturbation from the relationship between the visual feedback and reach angle, two variables that are entirely decoupled in the clamp condition. Nonetheless, we can simulate the CPC model with delayed contingent feedback. When tested with the designs used in Exps. 3-4, the model failed to predict the observed between- and within-block features of learning in those conditions (See Supplementary Result 3, Figs. S3-4).

## Discussion

In the current study, we identified a novel form of implicit learning, what we refer to as implicit aiming in the context of sensorimotor adaptation. Over a series of experiments and simulations, we found that, in contrast to implicit recalibration, implicit aiming is less constrained by feedback delays, exhibits savings-in-relearning, and follows unique temporal dynamics (See Table 1 for a summary). These characteristics closely resemble those associated with explicit aiming, suggesting that they reflect features of processes involved in action selection rather than action execution. These findings are consistent with our proposed alternative that the taxonomy of the processes involved in sensorimotor learning should be organized along computational goals associated with action selection and movement execution, rather than along the phenomenological distinction between implicit and explicit processes (Fig. 1).

**Table 1:** Different characteristics of implicit aiming and implicit recalibration.

|  | Implicit Aiming | Implicit Recalibration |
| --- | --- | --- |
| Sensitivity to feedback delay | Retained under delayed feedback | Suppressed by delayed feedback |
| Automaticity | Requires an intention to learn | Operates without intention |
| Dynamics in relearning | Savings in relearning | Attenuation in relearning |
| Contextual sensitivity | Modulated by context | Insensitive to context |
| Potential computational model | COIN | CPC |

### An implicit contribution to action selection

It is important to articulate what we mean by ‘implicit’ in the present work. Here we have adopted an operational definition that has been employed in many studies of sensorimotor adaptation^7,9,13,25^. Namely, we refer to a learning process as implicit when a consequence of its operation persists in an aftereffect phase in which participants have been informed that the perturbation has been removed and are explicitly instructed to reach directly towards the target. Perhaps the strongest evidence for the existence of an implicit aiming comes from Exp. 3 where we used delayed feedback to suppress implicit recalibration and limited planning time to suppress explicit aiming. Despite these validated, powerful, and popular techniques to suppress learning, we still observed prominent learning.

Unlike implicit recalibration, implicit aiming appears to be influenced by cognitive factors. We observed no adaptation in the delayed clamp condition (Exp. 1 control), suggesting that implicit aiming only operates when there is an intention to learn. Moreover, we showed that this process is clearly influenced by how the task is framed (Exp. 2). Priming participants to anticipate that error is due to an external perturbation decreased implicit aiming whereas priming participants to expect error from endogenous movement biases increased implicit aiming. This flexibility is not observed with implicit recalibration: Here we see an automatic response to sensory prediction error, even when this response is maladaptive in terms of task success^8,56^.

The priming results provide insight into the nature of implicit aiming. We hypothesize that implicit aiming arises as part of an implicit credit assignment problem. In response to the error arising from a perturbation, participants may consciously adopt an aiming strategy (i.e., explicit aiming). But there is likely uncertainty associated with the source of this error: How much is due to a perturbation and how much is due to internal biases that have emerged over the course of learning? When instructed to stop aiming, we assume that the participant can immediately nullify an aiming strategy. The residual error comprises implicit aiming, the implicit change in behavior designed to nullify biases introduced by the sensorimotor system. By manipulating the is credit assignment process, we were able to influence the relative contribution of implicit aiming and explicit aiming.

There remain a number of open questions regarding the nature of implicit aiming. First, the degree to which implicit aiming operates in parallel or in tandem with explicit aiming remains to be determined. One possibility is that implicit aiming and explicit aiming might be two processes built up separately during training. Alternatively, there might be one aiming process during training, with the assignment to external vs. internal sources only becoming relevant when participants were asked to move directly to the target in the aftereffect phase. Second, it is possible that there are multiple forms of implicit aiming, and that these other forms may not arise from uncertainty in credit assignment. We observed that, when preparation time was limited (Exp. 3a), learning was predominantly based on implicit aiming. Requiring fast RTs should increases motor noise. As such, the system would be more likely to attribute the error to an internal source, resulting in larger implicit aiming^11,48^. It is also possible that action selection might become automatized under the pressure of limited RT^57^, making it difficult to eliminate this in the washout phase. However, the time-pressure constraints in Exp3, which prevented explicit aiming, do not support the hypothesis that explicit strategies automatize into implicit learning.

### Distinct properties between implicit aiming and implicit recalibration

Having identified a signature of implicit aiming in Exps. 1 and 2, we went on to explore the features of this process. The results of Exps. 3 and 4 highlighted how implicit aiming and implicit recalibration are strikingly different in terms of their temporal dynamics. Importantly, these distinctions help reconcile previously observed discrepancies in the literature^13,14,22,24,25,46^.

One major controversy in the motor learning literature concerns how implicit learning processes operate in a savings paradigm. Experimentally, some studies have shown that the implicit system exhibits savings, whereas others have reported that relearning is actually slower than initial learning^13,23–25,46^, results that were hard to reconcile if one assumes that the same implicit process is being probed in these studies. The results of Exp. 3 provide a resolution to this controversy, indicating that implicit aiming exhibits savings in relearning while implicit recalibration exhibits anti-savings^24^. We propose that the opposing results reported in previous studies reflect differing contributions from implicit aiming and implicit recalibration. For instance, in studies involving small perturbations^13^ or limited reaction times^25^, implicit aiming is likely to be relevant, and thus one will find savings. In contrast, studies using clamped feedback isolate implicit recalibration find anti-savings because the recalibration process is subject to interference from recent experience^23,24^.

A second controversy has centered on the temporal dynamics of implicit adaptation given evidence that this form of learning exhibits both volatile and stable components. When clamped feedback is used to isolate implicit recalibration, the volatile component appears to be modest, typically contributing no more than ∼4° of change^22,23^. In contrast, studies measuring implicit learning with typical rotational feedback have reported that the volatile process may contribute up to 10° of learning^13^, at least in the early phases of training. We hypothesize that the measurement of implicit learning in the latter case reflects contributions from both implicit aiming and implicit recalibration. Indeed, when we measured the temporal dynamics of the two implicit components separately in Exp. 4, both showed a change in hand angle after the 1-min break and their combined magnitude was close to what has been observed in previous works^13^.

Moreover, we propose that the reduction in hand angle after the break arises from the different mechanisms associated with implicit recalibration and implicit aiming. For implicit recalibration, the drop of hand angle after each 1-min break reflects decay of a volatile memory; for implicit aiming, the drop stems from increased uncertainty in contextual inference rather than forgetting per se. We assume that for implicit aiming, training trials have a cumulative effect to strengthen the representation of the perturbed context whereas extended baseline trials strengthen the representation of the baseline context. Consistent with this assumption, the magnitude of the post-break drop decreased across training (Exp. 4) but increased when participants experienced a long baseline phase before training (Exp. S3). In contrast, neither the length of the baseline or the training session had any influence on the stability of implicit recalibration memory, suggesting the volatile component here is not due to contextual uncertainty.

### New taxonomy for sensorimotor learning

Harkening back to the seminal studies of medial temporal lobe amnesia, taxonomies of learning have tended to be organized on a phenomenological level, featuring a primary branch between explicit and implicit processes^4,58–60^. This organization has been adopted to describe the multiple learning processes contributing to sensorimotor adaptation.^2,7^ Over the years, however, a number of ambiguous findings has made it increasingly difficult to place their associated processes squarely under the distinct branches of this explicit-implicit taxonomy. The current results offer a different organizational principle, distinguishing between learning processes based on their underlying computational goals, to reconcile this problem. As shown in Fig. 1, the top branch distinguishes between processes involved in action selection and processes involved in action execution. The former chooses the optimal action to achieve a desired goal, whereas the latter is designed to ensure that the selected movement is executed with optimal accuracy and efficiency.

The key computational differences between action selection and action execution are captured by the COIN and CPC models, respectively. The COIN model involves representations of multiple motor memories associated with different contexts and allows for flexible switching between these memories based on inferred context^61^. This mechanism allows the action selection system to rapidly implement appropriate behaviors in response to varying environmental demands; for example, switching from controlling a cursor with a mouse to using a trackpad. In contrast, the CPC model does not represent discrete contexts. Instead, learning reflects a continuous recalibration based on the previous state. This aligns with the idea that this system is designed to automatically correct for internal changes in the motor system itself (e.g., fatigue, proprioceptive drift), changes that tend to be gradual.

This new taxonomy offers a parsimonious framework that reconciles past contradictions and opens new avenues for investigating the multifaceted nature of motor learning. Although we have focused on sensorimotor adaptation in this paper, we believe this revised taxonomy will be applicable to motor learning in general. For example, a skilled baseball pitcher will draw on their explicit knowledge of the opponent’s hitting strengths and weaknesses to choose a type of pitch and desired location. However, this plan may be altered by the implicit knowledge gained from past encounters with this opponent. The final outcome of the encounter will also, of course, depend on how well calibrated the execution system is to transform the action command into a movement. Beyond aiming and recalibration, the new taxonomy can be expanded to specify the computational characteristics of other learning processes. For example, habituation and reward-related learning could be attributed to action selection, while lower-level processes, such as efficient coding of movement direction would fall under the action execution branch. A taxonomy built upon the distinction between action selection and action execution may also prove useful in considering the shift in control across cognitive and autonomous stages during skill acquisition.

## Method

### Participants

For lab-based experiments, we recruited a total of 188 undergraduate students from the University of California, Berkeley (106 female, mean age = 21.5, SD = 3.6). For web-based experiments, we recruited 91 participants (46 female, mean age = 28.1, SD = 4.5) from Prolific.co. We aimed to include approximately 20 participants per condition in the in-person experiments and around 30 participants per condition in the online experiments, consistent with typical sample sizes in motor adaptation studies. All participants were right-handed, as determined by their scores on the Edinburgh Handedness Inventory^62^ and had normal or corrected-to-normal vision. Participants were compensated at a rate of $20 per hour. Participant demographics for each experiment are summarized in Table S1. All experimental protocols were approved by the Institutional Review Board at the University of California, Berkeley.

### Apparatus

Lab-based experiments: Participants performed a center-out reaching task on a digitizing tablet (Wacom Co., Kazo, Japan) which recorded the position of a digitizing pen held in the hand. Stimuli were displayed on a 120 Hz, 17-inches monitor (Planar Systems, Hillsboro, OR) mounted horizontally above the tablet and, thus, occluding vision of the arm. The experiment was controlled by a Dell OptiPlex 7040 computer (Dell, Round Rock, TX) running on a Windows 7 operating system (Microsoft Co., Redmond, WA) with custom software coded in MATLAB (The MathWorks, Natick, MA) using Psychtoolbox extensions^63^.

Web-based experiments: The code was written in JavaScript and presented via Google Chrome, designed to run on any laptop computer. Visual stimuli were presented on the participant’s laptop monitor and movements were produced on the trackpad. Data was collected and stored using Google Firebase.

### Procedure

#### Experiment 1

Implicit aiming task: We designed a modified version of a visuomotor rotation task to measure implicit aiming. The start position (radius: 4 mm) was located at the center of the screen, and a single blue target (radius: 8 mm) was positioned at 60° (0° = rightward, 90°= outward), 12 cm from the center. This target location was chosen based on previous research showing that there is minimal directional bias for reaches to this location (Wang et al. 2024). Participants were instructed to execute a rapid slicing movement to intersect the target. Upon reaching the target distance, a tone was played to signal that they had moved far enough.

In trials with cursor feedback, the cursor (radius: 3 mm) appeared 1.5 seconds after this tone. The radial distance to the cursor from the center was always 12 cm, the target distance. The rule determining the angular position varied across the experiment (see below). We delayed the feedback by 1.5 s to eliminate implicit recalibration^11,37^. The cursor remained visible for 100 ms and then disappeared. In trials without cursor feedback, the trial ended when the tone was played (reach distance passed the target radius). If the movement duration exceeded 500 ms, an audio clip saying “Too Slow” played before the next trial.

Participants were instructed return to the start position after completing the reach. The cursor was blanked during the return movement and only reappeared when the hand was within 2 cm of the center. The position of the cursor during this return phase was always veridical with hand position. After maintaining at the start position for 500 ms, the blue target reappeared, prompting the next movement.

There was a total of 407 trials in Experiment 1. It began with a no-feedback baseline block (20 trials) followed by a baseline block with veridical feedback (20 trials). To ensure the participants could flexibly use an aiming strategy, we included two short blocks with a 65° rotation, one in the clockwise direction and one in the counterclockwise directions (20 trials per block). Participants were informed of the perturbation and instructed to re-aim in the opposite direction so as to make the cursor hit the target. Each rotation block was followed by a no-feedback block (15 trials) and then a feedback block (10 trials). Before each no-feedback block, the participant was instructed to stop aiming and move directly to the target. Participants showed minimal aftereffects in these two no-feedback blocks indicating that they understood the instructions and were able to rapidly terminate their use of an explicit aiming strategy.

The main part of the experiment consists of three rotation blocks in which the perturbation size was gradually changed: Block 1: 0 to 25°; Block 2: 25 to 65°; Block 3: 65° back to 25°. The rate of change within each block was 0.5° per trial. Each of the three blocks was followed by a no-feedback block (15 trials). Prior to each of the no-feedback blocks, the instructions were repeated, informing the participants that there would be no feedback cursor and that they were to reach straight to the target, dispensing with any aiming strategy. These no-feedback blocks provided our assay of implicit aiming.

#### Experiment S1

Exp S1 was to verify that the 1.5 s feedback delay effectively suppresses implicit recalibration. In this experiment, we used task-irrelevant clamped feedback on the perturbation trials. On these trials, the feedback cursor was again displayed at the target distance after a delay of 1.5 s, but the angular position was predetermined, independent of the participant’s hand position. The magnitude of the clamp matched the rotation size used in Exp 1 (i.e., 65° in the pre-experiment phase and ramped in the main experiment). Importantly, before each clamp-feedback block, participants were informed of the nature of the clamp feedback and instructed to ignore the feedback, reaching straight toward the target.

#### Experiment S2

Exp S2 was designed to test whether the aftereffects observed in Exp 1 could be attributed to use-dependent learning. The experiment began with a 60-trial no-feedback baseline block in which three targets were presented (30°, 50°, 70°). This was followed by 183 trials in which the target was always presented at 50°. The feedback was veridical for the first 10 trials and then following a gradual rotation for the next 173 trials, increasing linearly from 0 to 20° (0.15°/trial) before decreasing from 20° to 15° (0.15°/trial). The rotation block was followed by a short no-feedback block (5 trials), again with the target always at 50° target. The final block was composed of 40 no-feedback trials with targets 30° and 70° (20 trials/target) to assess generalization.

#### Experiment 2

Experiment 2 tested the hypothesis that implicit aiming arises from a credit assignment problem between errors attributed to external environmental changes or internal motor bias/drift. To this end, we used the basic design of Experiment 1, but manipulated the instructions across groups. We primed participants in the external bias condition to attribute the error to an external perturbation with the following instructions: “We will gradually introduce a rotational perturbation in the next session. Specifically, the rotation will change roughly 0.5° after each movement, increasing from 0° to 75°. After that, the rotation will decrease from 75° to 25°. Please adjust your movement accordingly to make the white dot appear on the target.”

For the participants in the internal bias condition, we provided the following instructions prior to the start of the gradual rotation block: “The cursor may or may not follow your movement in the next session. However, research has shown that one’s perception of the hand may gradually drift when you are performing repeated movement without seeing their hand. Please try your best to adjust your movement based on the white dot to make the cursor appear on the target.” In addition, we eliminated the two abrupt rotation blocks given that these would have highlighted an external perturbation. The remaining task procedure was the same as Experiment 1.

#### Experiment 3

Experiment 3 used a between-group design to compare the effect of implicit aiming and implicit recalibration on relearning. For both groups, we suppressed explicit aiming by using used an RT countdown procedure that minimizes the time available to plan the movement. After positioning the cursor in the start position, four 50-ms tones (each lasted 50 ms) were played with a fixed stimulus-onset asynchrony of 500 ms. The target appeared at one of three positions (70°, 190°, and 310°), 250 ms after with the third tone. The participant was instructed to use the auditory rhythm to time their movement to begin with the fourth tone. Thus, if they achieve this goal, there would only be 250 ms between the target onset and movement onset. To push participants to heed these instructions, the message “too slow” was displayed if the movement started more than 250 ms after the target onset (i.e., after the onset of the last tone).

For the implicit aiming group, the feedback was delayed by 1.5 to eliminated implicit recalibration. For the first learning phase, participants completed a 30-trial no-feedback baseline block, a 60-trial feedback baseline block, a 180-trial rotation block with a fixed 30° rotation, and a 30-trial no-feedback block to assess the aftereffect. This was followed by a 30-trial feedback block to washout learning from the first learning phase. For the second learning phase, participants completed a 30-trial no-feedback block followed by a second abrupt rotation block of 180 trials. Relearning is assessed by comparing performance in the second rotation block compared to the first rotation block.

For the implicit recalibration group, the rotational feedback was replaced by clamped feedback and the feedback appeared as soon as the amplitude of the hand movement reached the target distance (no delay). Participants were instructed to ignore the clamped feedback and aim directly at the target. The first learning phase was identical in structure to that used with the implicit aiming group: 30-trials no-feedback baseline block, 60-trials 0° clamp block, a 180-trial 30° clamp block (180 trials), and a 30-trial no-feedback block. To washout learning in the first block, we reversed the direction of the clamp (-30°) for 120 trials. The length of this block was based on the results of pilot work to determine when implicit recalibration from the initial 30° clamp block was fully eliminated. For the second learning phase, participants completed a second 30° clamp block of 180 trials.

#### Experiment 4

Exp 4 was designed to examine the temporal stability of implicit aiming and implicit recalibration. As in the other experiments, the feedback was delayed by 1.5 s for the implicit aiming group. The experiment began with a no-feedback baseline block (20 trials) followed by a rotation block (80 trials). The rotation size was initially set to 30° and then gradually decreased to 15° over the course of the block following an exponential decay. This perturbation schedule was designed to approximate the time course of errors that were experienced with a visuomotor rotation task in Hadjiosif et al.^13^ Part of the error in the classic visuomotor rotation task is compensated by implicit recalibration; thus, the error signal relevant for the aiming system decreased across time. To examine the temporal stability of the learning, we inserted three 60 s-breaks after trials 40, 70, and 100. A no-feedback washout block (20 trials) was applied after the rotation block to measure implicit aiming. For the implicit recalibration group, the delayed rotational feedback was replaced by non-delayed clamped feedback and participants was instructed to ignore the feedback.

#### Experiment S3

ExpS3 was designed to examine how prior experience with non-rotated feedback influences the temporal stability of implicit aiming and implicit recalibration. The overall design mirrored that of Exp 4, with one key difference: Following the initial no-feedback baseline, we introduced an additional 70-trial baseline block. We applied veridical feedback in the delayed feedback condition and 0° clamped feedback in the clamp condition.

### Analysis

The initial data analyses were conducted in MATLAB 2022b. Reach angle was calculated as the angular difference between the target and the hand position at the target radius. Positive values indicate reach angles in the opposite direction of the perturbation, the direction that would be expected by adaptation. Reaction time was defined as the first sample in which the hand surpassed the radius of the start position. Movement time was defined as first sample in which the radial distance of the hand position was great than the target distance, minus the reaction time.

To minimize online corrections, we instructed the participant to move quickly. Trials in which the movement time was longer than 1000 ms were excluded from the analysis (<0.5%). We also excluded as outliers, trials in which the cursor or reach angle (when clamp feedback was applied) at the end of the movement was more than 70° from the target under the assumption that the participant moved to the wrong target on these trials (<0.5%). For Exp 3, we removed trials with reaction time longer shorter than 0 ms or longer than 350ms (∼5%).

To calculate the retention rate in Exp 3 and 4, we fitted the average hand time course during the washout phase using a state-space model:

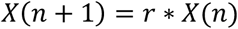

Where *X* is the hand angle, *n* is trial number, and *r* is the retention rate. We used a bootstrapping procedure, resampling with 1000 iterations, to estimate the standard deviation of the forgetting rate at the group level.

Aftereffects were defined as the mean reach angle over the no-feedback block following the perturbation, minus the mean of the first no-feedback baseline block. One sample t-tests were applied to examine whether the aftereffect is different from zero. Between-condition comparisons were performed with t-tests, with false discovery rate (FDR) corrections for multiple comparisons applied when appropriate. To analyze (anti)savings from the learning curves in Exps 4 and 6, we used a cluster-based permutation test, a method useful for data with temporal dependencies^24,48,49^. In all tests, we confirmed that the data met the assumptions of a Gaussian distribution and homoscedasticity.

## Models

### Contextual inference (COIN) model

The COIN model is a Bayesian framework that models sensorimotor learning as the inference of latent contexts based on observed motor errors.^49^ Rather than representing a single adaptive process, the model posits that learning involves dynamically assigning credit to multiple internal models, each associated with a different context. Contexts are inferred based on the statistical structure of sensory feedback and the history of actions and outcomes. On each trial, the learner updates both the probability of each context and the associated motor memory, balancing prior beliefs with sensory evidence.

We used the code and parameter settings described in Heald et al.^49^ to simulate the delayed feedback conditions in Exp 3-4 using the basic version of the COIN. To simulate the change in behavior after each 1-min break (Exp 4), we modeled the 1-min break as 15 no-feedback trials; thus, uncertainty will gradual increase given that participants receive no input concerning the current state.

Central to the COIN model is the assumption that participants make an inference about the context and its associated perturbation from the visual feedback and reach angle. However, in the clamp task, the feedback is not contingent on the reach angle. As such, the basic COIN model cannot be used to generate reasonable prediction with clamp feedback. We note that there are different ways to modify the basic model to use COIN to simulate the clamp task. First, we might assume that, given the instruction to ignore the feedback, the system makes no inference from the clamp feedback and thus generates no adaptation.

Second, the system might treat the feedback as if it is contingent feedback. However, since the change in reach angle does not reduce the error, the inferred perturbation size would be unbounded (may go to infinity). Third, we may assume that the system inferring the clamp size regardless of the reach angle. Under this assumption, the model will predict the same outcome for clamped feedback and delayed contingent feedback. Given that each of these approaches is problematic, we opted to not use clamped feedback in our COIN simulations.

### Cerebellar Population Coding (CPC) model

The CPC model was developed to account for the learning dynamics of implicit recalibration based on a cerebellar-like network. The formalization of the model is reported in our previous paper^23^. The model includes a basis set of directionally tuned units in the cerebellar cortex, and a downstream integrator in the DCN that generates motor output. Learning occurs through error-driven plasticity, specifically long-term depression (LTD) at parallel fiber–Purkinje cell synapses and long-term potentiation (LTP) at mossy fiber–DCN synapses. These synaptic changes alter the balance of activity across populations, allowing the system to adjust motor commands in response to persistent sensory prediction errors.

Here we used the CPC model to model the anti-saving effects observed for the clamped feedback group in Exp 3. The CPC model includes two learning sites, one at the level of Purkinje cells (PC) in the cerebellar cortex and a second at the level of the deep cerebellar nuclei (DCN). By setting different learning/retention rates for these two sites, we used the CPC to simulate a dual-rate model to evaluate the dynamics of implicit recalibration in Exp 3 and 4. We simulated the CPC model with the parameter values determined in the original development of the model^23^.

## Data and code availability

Data and analysis code will be made available upon publication.

## Author contributions

T.W., T.L.., J.T., R.B.I. contributed to the conceptual development of this project. T.L. collected the data. T.W. and T.L. analyzed the data, prepared the figures, and wrote the initial draft of the paper, with all the authors involved in the editing process.

## Acknowledgments

RBI was funded by grants NS116883 and NS105839 from the National Institutes of Health (NIH). JAT was funded by grants NS131552 and NS134754 from National Institutes of Health (NIH).

## Competing interests

RI is a co-founder with equity in Magnetic Tides, Inc.

## Supplementary Results

### Supplementary Result 1: 1.5s delayed feedback sufficiently suppressed implicit recalibration

We conducted a control experiment (Exp S1) to confirm that the aftereffect observed in Exp 1 was not due to implicit recalibration. For this experiment, we used the same perturbation schedule but replaced the hand-contingent feedback with task-irrelevant feedback (“error clamp”) and instructed participants to always reach directly to the target. If the cursor feedback is not delayed, participants gradually change their reaching direction without awareness in response to clamped feedback^8,56,64^. However, we delayed the clamped feedback by 1.5 s in this control condition, a manipulation that effectively nullifies implicit recalibration^11,37^. Consistent with this assumption, we did not observe any change in movement direction during the perturbation, nor was there a measurable aftereffect in the no-feedback blocks (Fig 2A). By inference, the absence of an aftereffect in this control experiment indicates that the aftereffects observed in the main experiment reflects the operation of a distinct implicit process. Indeed, a direct comparison of the aftereffect data showed that the group receiving contingent (task-relevant) feedback exhibited larger aftereffects compared to the control group (t(36)s>2.7, ps<0.009). Interestingly, the absence of an aftereffect in the delayed clamp condition suggests that implicit aiming does not operate when participants have no intention to learn.

### Supplementary Result 2: Implicit aiming is not caused by use-dependent learning

We have hypothesized that the aftereffect observed following the gradual perturbations in Exp 1 reflects the operation of an implicit process contributing to action selection. However, it is possible that the effect is a manifestation of use-dependent learning, a process whereby a movement is biased in the direction of recently repeated movements.^18,65^ The magnitude of use-dependent learning in visuomotor adaptation studies is around 2-5°^19,66^, similar to the aftereffect values observed in the two probe blocks following the abrupt, 65° rotation. However, the aftereffect values in three probe blocks during the gradual rotation phase were much larger (>10°), suggesting they are unlikely to be caused by use-dependent learning.

Nevertheless, we conducted a control experiment (Exp S2) to ask if these aftereffects were due to use-dependent learning. For this experiment, we ramped up the perturbation to 20° and then ramped it back down to 15° before a no-feedback block to assess the aftereffect. During this no-feedback block, the first five trials participants went to the training location. In the next 40 trials, the target randomly appeared at two alternative positions that flanked the target position by 20° to assess generalization (Fig S1B). If the aftereffect is due to use-dependent learning, we should observe a bias towards the training target; as such, the aftereffect will be in opposite directions for the two displaced targets. In contrast, if the aftereffect is due to implicit aiming, we would expect the bias will be in the same direction for all three targets.

Participants tracked the gradual rotation during the perturbation phase and, consistent with Exp 1, there was a significant aftereffect during the no-feedback block. Importantly, the aftereffect was in the same direction for all three targets, with the magnitude reduced or the two generalization targets. These data are consistent with the hypothesis that the aftereffect is due to an implicit change in aiming rather than a manifestation of use-dependent learning.

### Supplementary Result 3: The CPC model failed to predict the learning dynamics with delayed feedback

We simulated the CPC model for the delayed feedback conditions in Exp 3 and 4. For Exp 3, the CPC prediction was nearly identical to that of the clamp condition: The model again predicted an anti-savings effect rather than a savings effect (Fig S3A), at odds with what was observed in the implicit aiming condition of Exp 3. For Exp 4, the CPC model also predicted that the magnitude of the decrease would be reduced across the three probe blocks, similar to what was observed in the implicit aiming condition. This occurs because the contribution of the volatile process diminishes when the remaining error becomes smaller (Fig S3B). However, the magnitude of the predicted drop was significantly smaller than what was observed in the experiment.

To further compare the CPC and COIN models in predicting the temporal stability of implicit aiming, we conducted an additional experiment (Exp S3). This experiment replicated the design of Exp 4 but included a long baseline block with feedback before the perturbation training (Fig S4A). According to the COIN model, this extended block will strengthen the prior for the baseline context. As such, after each probe, participants will be more likely to infer that they are more likely to be in the baseline context and the drop in hand angle should be larger after the 1-min breaks compared to when there was no baseline as in the main experiment (Fig S4B). In contrast, the CPC model lacks contextual inference and therefore predicts an identical drop in both the long and no-baseline conditions.

Consistent with the COIN prediction, we found that including a baseline block increased the magnitude of the drop compared to the no-baseline condition of Exp 4 (Fig S4B, linear mixed model, t(162)=2.8, p=0.005). This effect cannot be captured by the CPC model. These findings provide additional evidence that the drop after the break in the delayed feedback condition reflects contextual uncertainty, consistent with the COIN model, rather than forgetting of a volatile process as posited by the CPC model. In contrast, for the clamped feedback condition, when 0° clamped feedback (i.e., cursor moves directly to target) was used in an extended baseline block (Fig S4C-D), the size of the drop remained the same as in the non-baseline condition (t(260)=0.62, p=0.56), consistent with the prediction of the CPC model.

**Fig S1.**
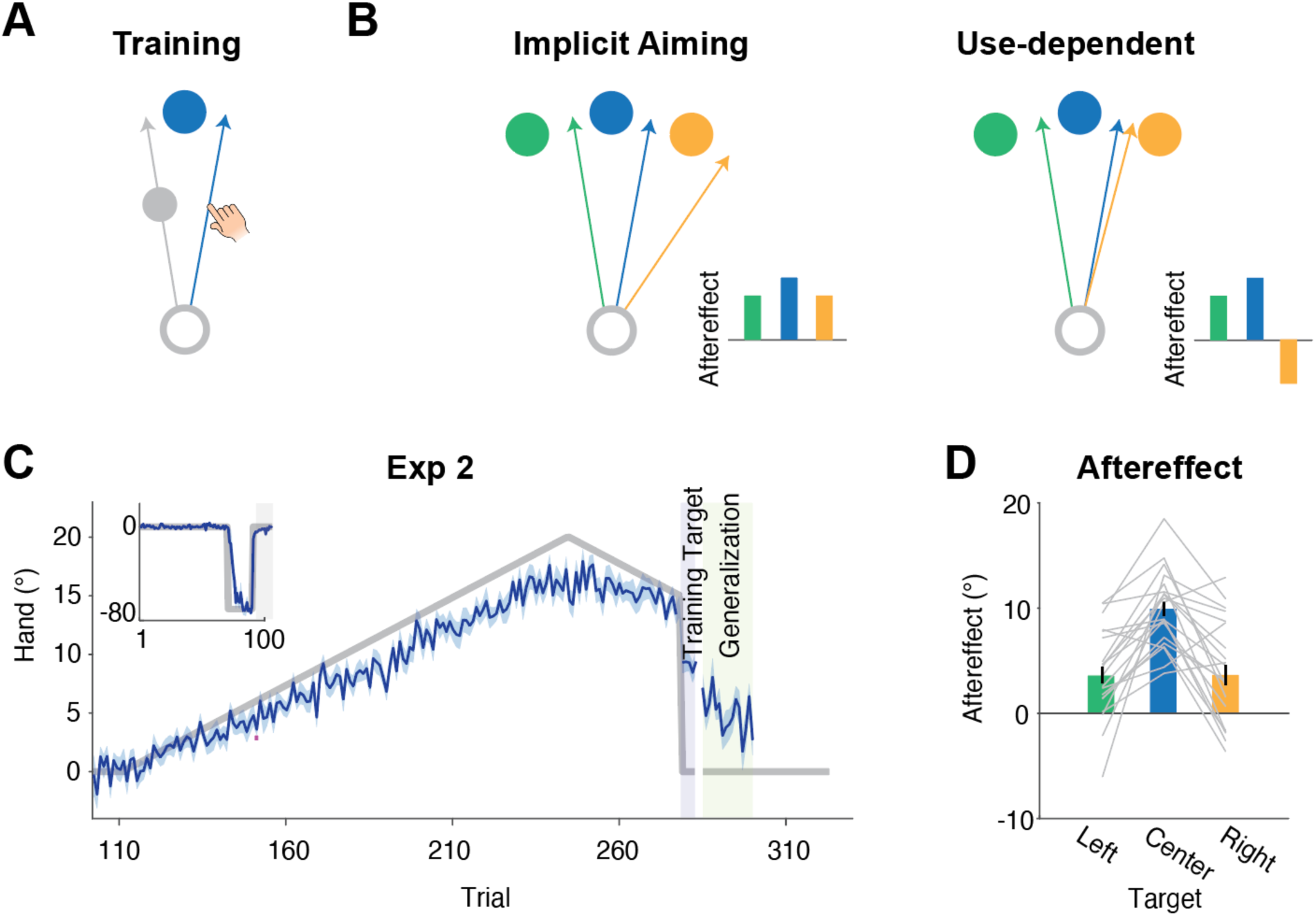
Residual aiming error observed during washout is not due to use-dependent learning. (A) The training target and schematic of hand path (blue) and rotated feedback (grey) late in the rotation block. (B) The target could appear at the training location or at one of two generalization locations in the no-feedback block, with the latter positioned 20° clockwise or counterclockwise from the training location. Generalization predictions for implicit aiming and use-dependent learning hypotheses. For the former, biases at the generalization locations should be in the same direction as for the training location. By the used-dependent hypothesis, biases at the generalization locations should be the opposite directions, both towards the training location. (C) Perturbation schedule (grey) and mean reach angle (blue). Only the training location was used in the first part of the no-feedback block (5 trials, light pink). In the second part of the block, two generalization locations were used (20 trials/target, yellow). (D) Reach angles in the no-feedback block are consistent with the pattern predicted by implicit aiming. Shaded areas in C and error bars in D indicate standard error.

**Fig S2.**
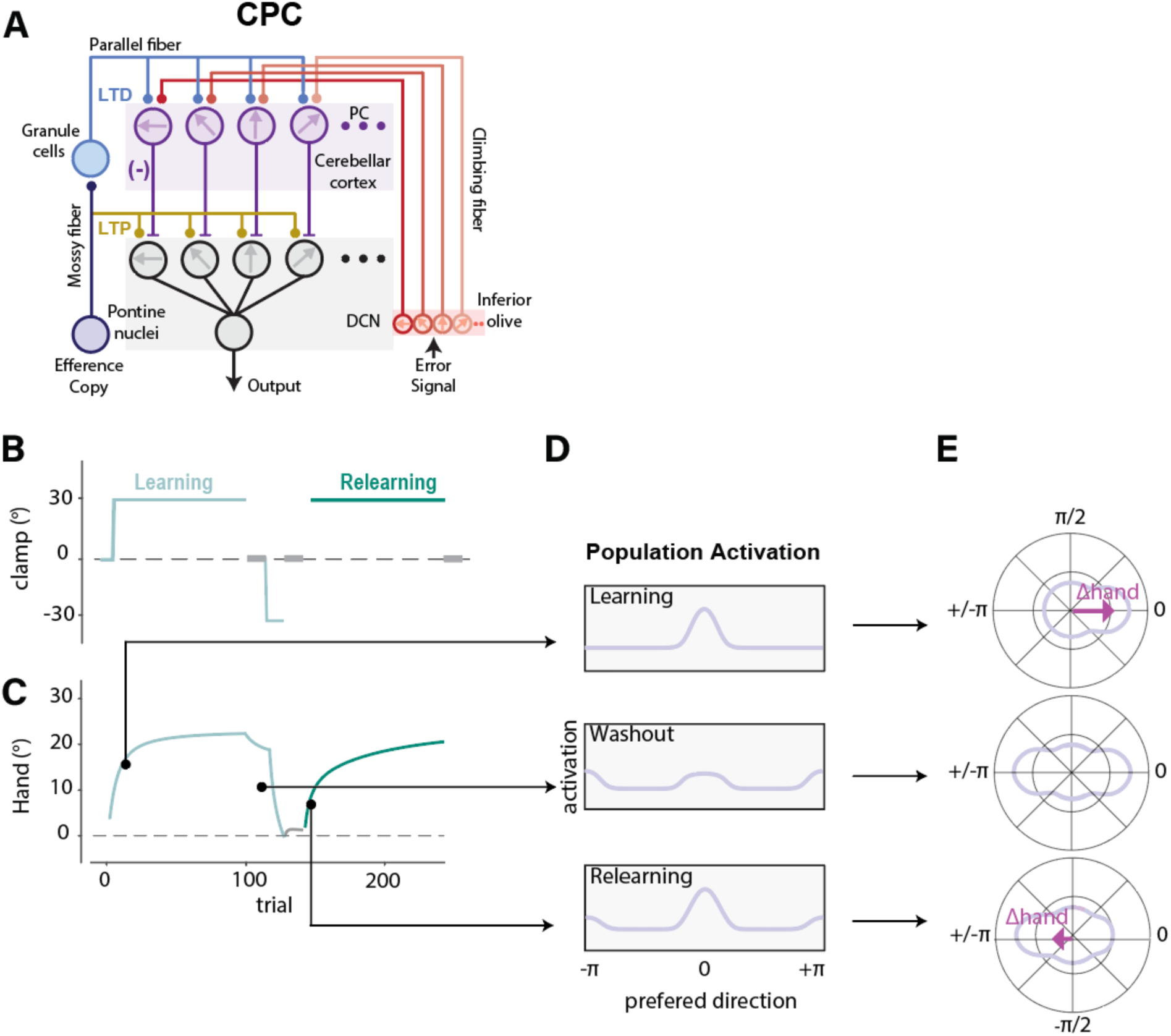
CPC model predicts an anti-saving effect. (A) Schematic of the CPC model. The CPC model assumes that learning occurs at both the PC layer and the DCN. Each layer includes a basis set tuned to different error directions. We have depicted only four units for simplicity. For a specific error signal, only the units tuned to that error direction will exhibit learning. (B) The perturbation schedule for Exp 3. (C) The predicted learning curve by the CPC model. (D) Population activation across the DCN basis set at three time points during the experiments. The x-axis indicates the preferred direction. The y-axis indicates the strength of activation. Top: In the first training block, the units tuned to the original perturbation were activated. Middle: However, during the washout, participants experienced opposite errors, and the units tuned to the opposite direction were also activated. Importantly, the forgetting of the previous learning is slow, so those units that were tuned to the original error directions are still activated, causing interference. Bottom: Similarly, in the second learning block, the units tuned to the opposite directions remain activated due to slow forgetting, which interferes with relearning and cause anti-saving. (E) Polar graphs showing how population activation translates into a change in hand angle. The angular axis indicates preferred direction, and the polar axis indicate strength of activation. The light purple line indicates the population activation, and the dark purple arrow represents the summation across all units, which determines the change in heading angle.

**Fig S3.**
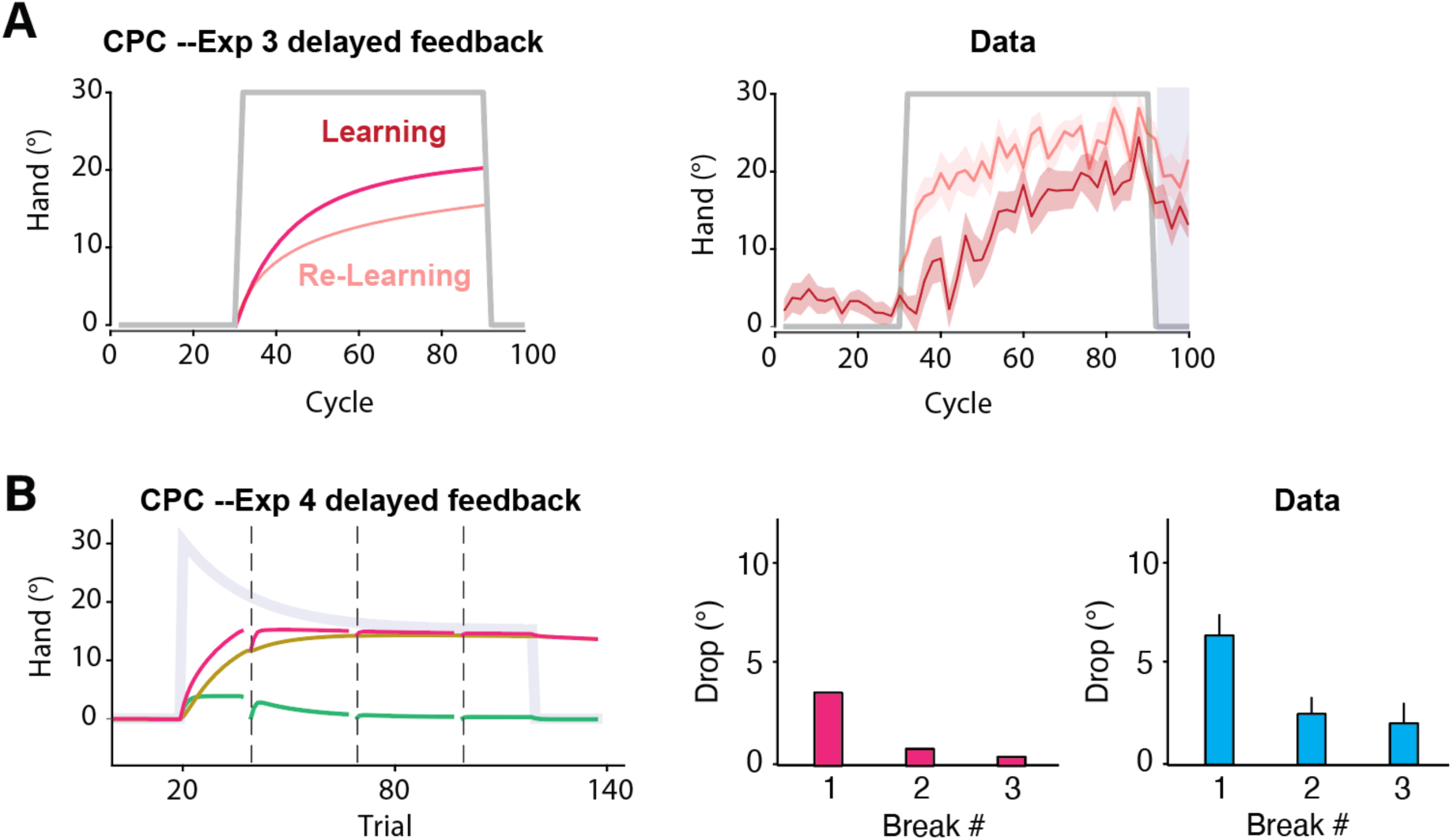
CPC model predictions for delayed feedback conditions. An important feature of implicit recalibration is that the error correction response does not scale linearly with error size ^23,67^. To capture this nonlinearity, we assumed that the error response increases linearly up to 7° and then saturates for larger errors ^8,56^. (A) Under this assumption, the CPC model generates nearly identical predictions for the delayed feedback condition in Exp 3 as for the clamped feedback condition, namely an anti-savings effect. This prediction is opposite to the savings observed in the behavioral results. (B) Similar to the observed results of Exp 4, the CPC model generates a reduction in the size of the drop since the contribution from the volatile process decreases across time with small errors. However, the magnitude of the predicted drops is substantially smaller than what was observed empirically.

**Fig S4.**
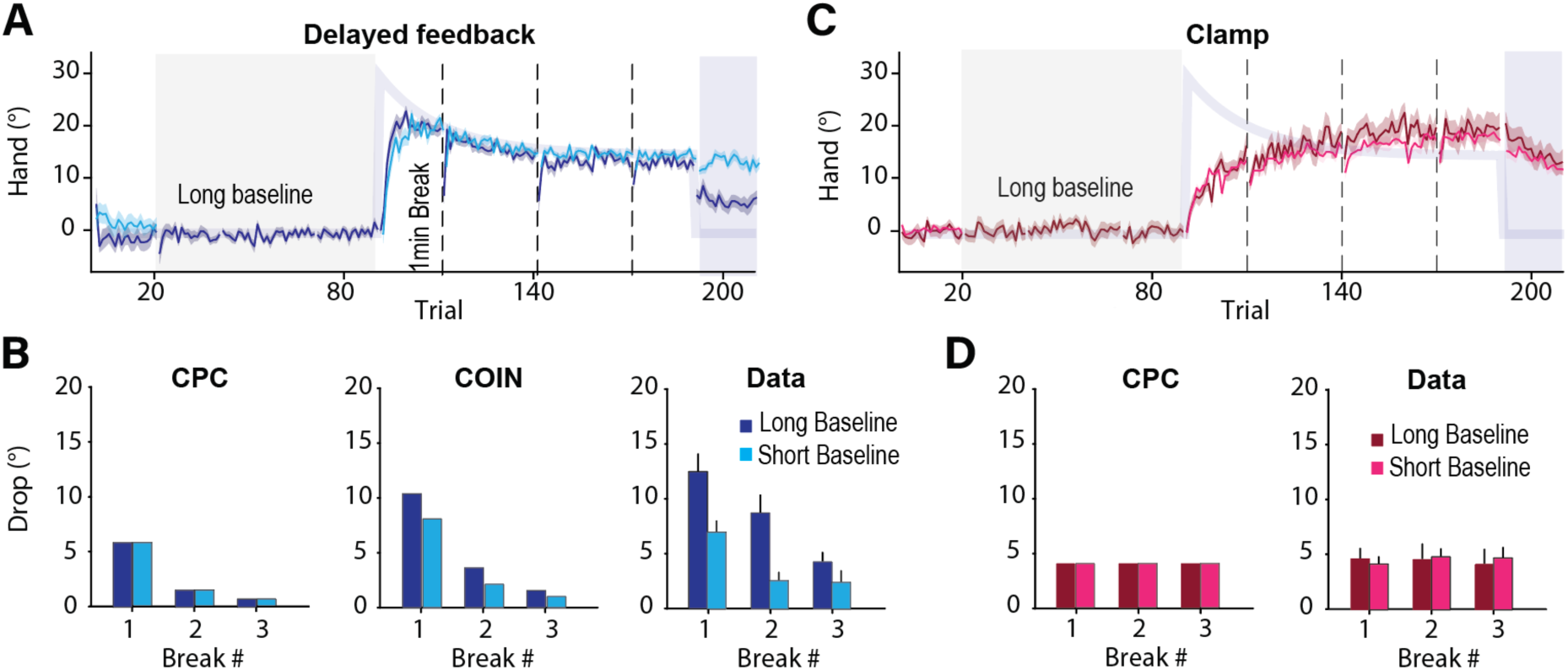
Differential effect of context on temporal properties of implicit aiming and implicit recalibration. (A) Time course of hand angle in the long (dark blue) and short baseline (light blue) conditions. Note that the break in the blue trace is for alignment purposes, enabling comparison of initial trials across phases. Vertical dashed lines indicate 1-minute breaks. (B) Model predictions for the CPC model (left) and COIN model (middle). Note that only the COIN model predicts a difference between the two baseline conditions (short is condition reported in main paper) across the three 1-minute breaks. Behavioral results (right) revealed an interaction between baseline length and the magnitude of the drop, consistent with the COIN model predictions but not the CPC model. (C) Time course of hand angle with non-delayed clamped feedback. (D) The CPC model correctly predicts that the drop in reach angle remains consistent across the three 1-minute breaks and is unaffected by the number of baseline trials.

**Table S1.**
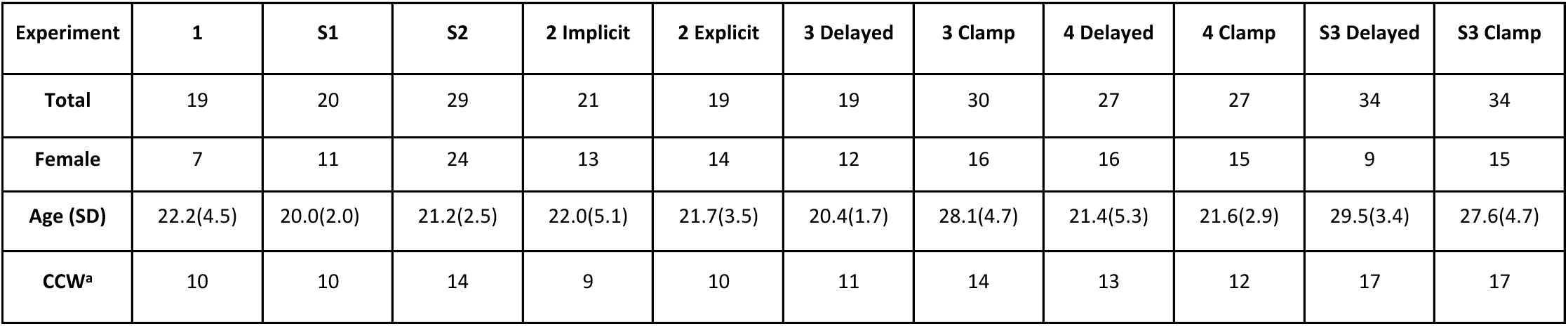
Participant Demographic information. ^a^: Number of participants who experienced a rotation in the counterclockwise direction during the main perturbation block. The other participants experienced a rotation in the clockwise direction.

| Experiment | 1 | S1 | S2 | 2 Implicit | 2 Explicit | 3 Delayed | 3 Clamp | 4 Delayed | 4 Clamp | S3 Delayed | S3 Clamp |
| --- | --- | --- | --- | --- | --- | --- | --- | --- | --- | --- | --- |
| Total | 19 | 20 | 29 | 21 | 19 | 19 | 30 | 27 | 27 | 34 | 34 |
| Female | 7 | 11 | 24 | 13 | 14 | 12 | 16 | 16 | 15 | 9 | 15 |
| Age (SD) | 22.2(4.5) | 20.0(2.0) | 21.2(2.5) | 22.0(5.1) | 21.7(3.5) | 20.4(1.7) | 28.1(4.7) | 21.4(5.3) | 21.6(2.9) | 29.5(3.4) | 27.6(4.7) |
| CCW<sup>a</sup> | 10 | 10 | 14 | 9 | 10 | 11 | 14 | 13 | 12 | 17 | 17 |

